# MHCflurry: open-source class I MHC binding affinity prediction

**DOI:** 10.1101/174243

**Authors:** Timothy O’Donnell, Alex Rubinsteyn, Maria Bonsack, Angelika Riemer, Jeff Hammerbacher

**Affiliations:** Department of Genetics and Genomic Sciences, Icahn School of Medicine at Mount Sinai, New York, New York, USA; Immunotherapy & Immunoprevention, German Cancer Research Center (DKFZ), Heidelberg, Germany; Molecular Vaccine Design, German Center for Infection Research (DZIF), partner site Heidelberg, Germany; Department of Microbiology and Immunology, Medical University of South Carolina, Charleston, South Carolina, USA

**Keywords:** MHC, HLA, epitope prediction, neural network

## Abstract

Machine learning prediction of the interaction between major histocompatibility complex I (MHC I) proteins and their small peptide ligands is important for vaccine design and other applications in adaptive immunity. We describe and benchmark a new open-source MHC I binding prediction package, MHCflurry. The software is a collection of allele-specific binding predictors incorporating a novel neural network architecture and adhering to software development best practices. MHCflurry outperformed the standard predictors NetMHC 4.0 and NetMHCpan 3.0 on a benchmark of mass spec-identified MHC ligands and showed competitive accuracy on a benchmark of affinity measurements. The accuracy improvement was due to substantially better prediction of non-9-mer peptide ligands, which offset a narrowly lower accuracy on 9-mers. MHCflurry was on average 8.6X faster than NetMHC and 44X faster than NetMHCpan; performance is further increased when a graphics processing unit (GPU) is available. MHCflurry is freely available to use, retrain, or extend, includes Python library and command line interfaces, and may be installed using standard package managers.

## Background

Adaptive immunity against intracellular infections and cancers depends on T cell recognition of small protein fragments (peptides) bound to major histocompatibility complex I (MHC I) proteins on cell surfaces. There are thousands of variants, or alleles, of MHC I proteins in the human population, each with specificity for binding a distinct set of peptides, which, when displayed by MHC, can potentially be the target of an immune response. Computational prediction of the binding affinity between a specified peptide and MHC allele has found wide application in infectious diseases, autoimmunity, vaccine design, cancer immunotherapy [1–4].

The NetMHC family of tools, which includes NetMHC[5] and NetMHCpan[6], are considered the state of the art predictive models for this task[7]. Both NetMHC and NetMHCpan are ensembles of shallow neural networks trained on binding affinity measurements deposited in the immune epitope database (IEDB)[8]. NetMHC uses an “allele-specific” approach, in which separate predictors are trained for each MHC allele; the input to the neural network is the peptide of interest. NetMHCpan uses a “pan-allele” approach, in which a single neural network takes as input both the peptide and a representation of the MHC allele. The impressive accuracy of these tools has resulted in wide adoption, despite certain limitations: they are closed source, may be trained (fit) only by their developers, and do not make available their training data.

Here we describe and benchmark a new package of allele-specific class I MHC binding predictors, MHCflurry version 0.9.1. MHCflurry predictors show competitive accuracy with the NetMHC tools and a significant speed improvement while addressing a number of limitations of the NetMHC software. In particular, MHCflurry is open source, retrainable, precisely documents the data and workflow used to train the released models on measurements in IEDB, exposes both a Python API and a command line interface, is installed using standard Python package management tools, and applies software development best practices such as unit tests, continuous integration, and code documentation.

MHCflurry is freely available under the Apache License 2.0. It may be installed from the Python package index. Source code is maintained at https://github.com/hammerlab/mhcflurry. All data and scripts used to train the models are available in this repository.

## Implementation

MHCflurry is implemented in Python (versions 2.7 and 3.4+ are supported) using the Keras neural network library (https://github.com/fchollet/keras). Model training and prediction use a graphics processing unit (GPU) when available.

Similar to NetMHC, MHCflurry is an ensemble of MHC I allele-specific predictors. Separate models are trained for each allele. The input to each model is a peptide. No representation of the MHC allele, such as its amino acid sequence, is used. The models are trained independently and no information is shared between alleles. For each allele, MHCflurry includes an ensemble of eight models. The final nanomolar affinity prediction is taken to be the geometric mean of the individual model outputs. The variance of the individual model predictions gives an indication of the uncertainty of the prediction and is also made available to users.

We arrived at the MHCflurry input representation, architecture, and training approach through an informal, iterative process using data held out from the training dataset. The final predictors were trained on the full training dataset and validated on the two benchmarks presented here.

### Peptide representation

MHCflurry introduces a novel peptide representation, in which variable-length peptides of length 8-15 are encoded as fixed-length vectors. Unlike the averaging scheme implemented in NetMHC and NetMHCpan, in which non-9mer peptides are encoded as 9-mers by adding or removing amino acids, this approach makes the full peptide available to the network. It also avoids the need for more complex neural network architectures, such as recurrent networks, that would be required to explicitly support variable length model inputs.

The motivation for the encoding is to preserve the positionality of the residues that make the most important stabilizing contacts with the MHC molecule. These are known as anchor positions, and generally occur toward the beginning or end of the peptide for most alleles. For example, the anchor positions of HLA-A*02:01 and many other alleles are at the second and last positions in the peptide, i.e. positions 2 and 9 for a 9-mer peptide. While overhangs, in which longer peptides protrude from the end of the binding groove, have been reported [9–12], it is thought that the most common conformation adopted by longer peptides is a bulged or zigzag conformation of the middle residues [13–15]. In this case the positions of the anchor residues remain intact relative to the nearest end of the peptide.

The peptide encoding is performed as follows (Figure 1a). Each peptide of length 8-15 is transformed to a length 15 string, in which missing residues are filled with an *X* character, which is treated as a 21st amino acid. The first four and last four residues in the peptide always map to the first four and last four positions in the representation. The middle seven residues are filled as needed depending on the length of the peptide: an 8-mer leaves all middle positions as an *X* whereas a 15-mer fills all positions. In this way, the positions most likely to contain anchor residues are consistently mapped to the same positions in the representation relative to the end of the peptide.

**Figure 1:**
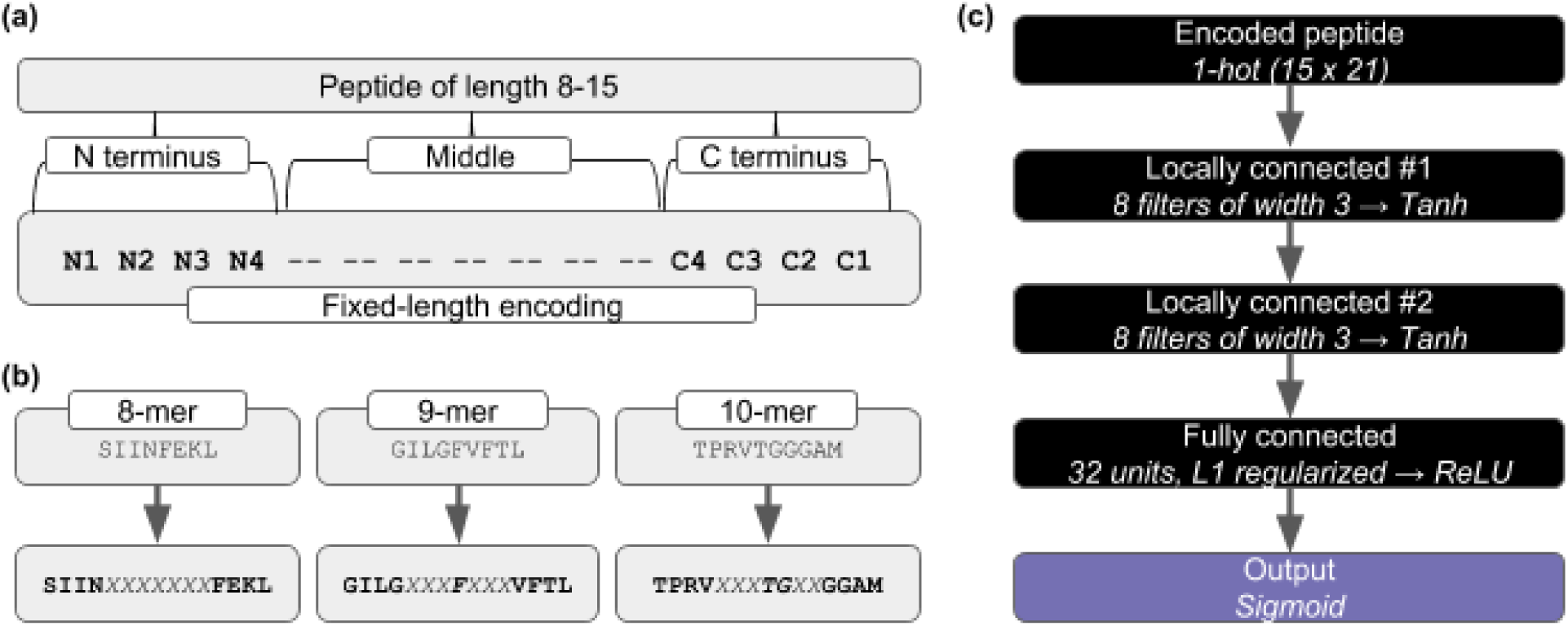
MHCflurry peptide representation and neural network architecture. **(a)** Variable length peptides (8-15mers) are encoded as length-15 sequences. The four N-terminal and four C-terminal residues map to fixed positions (N1-N4 and C1-C4) in this encoding. The seven middle residues are filled according to the length of the peptide, with unfilled positions set to the special character *X*. **(b)** Example encodings for three peptides. **(c)** Neural network architecture. The length-15 encoded peptide is supplied as a 1-hot vector, with entries for the 20 amino acids plus the *X* special character. Two locally connected layers are applied with a hyperbolic tangent (tanh) activation, followed by a fully connected layer with rectified linear (ReLU) activation, and an output neuron with sigmoidal activation.

### Output encoding

As in NetMHC and NetMHCpan, MHCflurry transforms binding affinities to values between 0.0 and 1.0, where 0.0 is a non-binder and 1.0 is a strong binder. The neural networks are trained using the transformed values and the inverse transform is used to return prediction results as nanomolar affinities. The transform is given by 1 - log_50000_(x) where x is the nanomolar affinity. Affinities are capped at 50,000 nM.

### Neural network architecture

The MHCflurry predictors are feedforward neural networks composed of the following layers: the peptide representation (described previously) encoded as a 1-hot (binary) vector, two locally connected layers, a fully connected layer, and a sigmoidal output (Figure 1b). Locally connected layers are one dimensional convolutional layers without weight sharing. Each neuron receives a neighborhood of adjacent points, instead of the full input from the previous layer as in a fully connected layer. The locally connected layers use hyperbolic tangent (tanh) activations, and the fully connected layer uses a rectified linear (ReLU) activation. The weights of the fully connected layer are L1 regularized.

In preliminary investigations (data not shown), we observed that models using more than one fully connected layer performed poorly and that using an amino acid embedding layer did not significantly outperform a 1-hot encoded peptide. We additionally tested two recent ideas from the deep learning literature, dropout[16] and batch normalization[17], which also did not significantly improve performance. While the downloadable MHCflurry models do not use these features, the MHCflurry software includes support for them to enable experimentation.

### Construction of the training dataset

The training dataset was assembled from a snapshot of IEDB MHC ligands downloaded on May 17, 2017 augmented with the BD2013 dataset published in ref. [18].

IEDB entries with non-class I, non-specific, mutant, or unparseable allele names were dropped, as were those with peptides containing post-translational modifications or noncanonical amino acids. To avoid the potential for bias in favor of MHCflurry on the mass-spec validation dataset, entries deriving from mass-spec studies were removed from the training data. This yielded an IEDB dataset of 147,716 quantitative and 43,704 qualitative affinity measurements. We assigned nanomolar affinities to the qualitative measurements as follows: *positive-high*, 50; *positive-intermediate*, 500; *positive-low*, 5000; *positive*, 100; *negative*, 50000.

Of 179,692 measurements in the BD2013 dataset published in ref. [18], 55,473 were not also present in the IEDB dataset. After selecting peptides of length 8-15 and dropping alleles with fewer than 200 measurements, the combined training dataset consists of 235,597 measurements across 101 alleles (Table S1).

### Neural network training

The MHCflurry models are trained (fit) using a procedure similar to NetMHC. Eight models are trained for each allele. Each model is trained on a random 80% sample of the data for the allele; the remaining 20% is used as a test set for early stopping. Training proceeds using the RMSprop optimizer[19] until the accuracy on the test set has not improved for ten epochs. Mean squared error is used as both the training loss and test set accuracy metric. At each epoch, 25 synthetic negative peptides for each length 8-15 are randomly generated. These random negative peptides are sampled so as to have the same amino acid distribution as the training peptides and are assigned uniformly random affinities between 20,000 nM and 50,000 nM. No model selection is performed.

The models within an ensemble thus differ from each other due to several sources of randomness. The most important factor is that each model is trained on a different 80% sample of the data. Lesser factors include random initial model weights, the nondeterminacy of stochastic gradient descent, and the random negative peptides.

Training the models described here took 311 minutes on a machine with eight 2.30 GHz Intel Xeon CPUs, one NVIDIA Tesla K80 GPU, and 52 GB memory.

## Benchmark approach

As the NetMHC tools are fit to binding data in IEDB, new datasets not included in IEDB are required to assess the performance of these tools. We use two datasets for this purpose: (1) a published dataset of peptides eluted from cell-surface MHC and identified by mass-spec[20], which we refer to as the ABELIN dataset, and (2) an unpublished dataset of affinity measurements generated through an HPV vaccine development project, referred to as the HPV dataset. All entries in both the ABELIN and HPV benchmarks are distinct from entries in the TRAIN dataset.

The ABELIN dataset was derived from 20,451 sequences of MHC-displayed ligands eluted from a B cell line and identified by mass spec by [20]. Each experiment was performed in cells engineered to express a single MHC I allele. We excluded 2/16 alleles (HLA-A*02:04 and HLA-A*02:07; Table S1) not supported by MHCflurry due to insufficient representation in the TRAIN dataset (fewer than 200 measurements) and discarded peptides with post-translational modifications or lengths outside the supported range (8-15 residues). To create the benchmark, we sampled unobserved sequences (decoys) from the protein-coding transcripts that contained the identified peptides (hits) based on protein sequences in the UCSC hg19 proteome [21] and transcript quantifications from RNA sequencing of the relevant cell line (b721221) downloaded from the Gene Expression Omnibus (accession GSE93315). For an allele with n hits, we sampled 100n decoys, weighting transcripts by the number of hits and sampling an equal number of decoys of each length 8-15. This yielded 2,045,100 decoys for 20,451 hits, from which we removed 118 (0.005%) entries also present in the TRAIN dataset, for a benchmark of 20,361 hits and 2,045,072 decoys. We assessed the accuracy of each predictor at differentiating hits from decoys in terms of positive predictive value (PPV). To compute PPV for an allele with n hits, we ranked the n + 100n hits and decoys from tightest to weakest predicted binding affinity and calculated the fraction of the top n peptides that were hits.

In addition to the standard MHCflurry, NetMHC, and NetMHCpan predictors, we considered seven variations on the MHCflurry architecture in the ABELIN benchmark. The variants changed one or two aspects of the architecture or training data and were otherwise identical to MHCflurry 0.9.1 (Table 1). For each architecture, we evaluated both a single predictor and an ensemble of eight models.

**Table 1:**
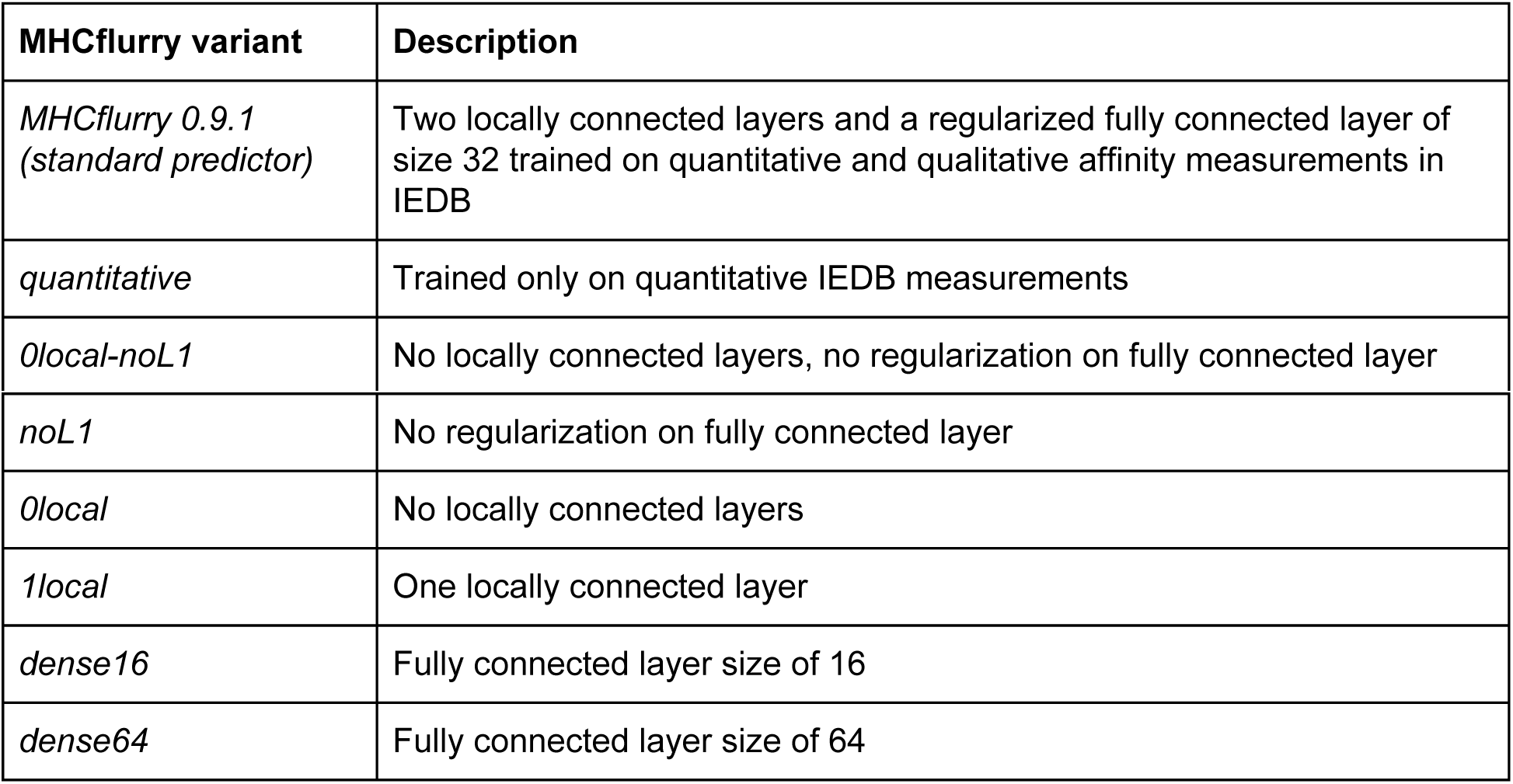
MHCflurry architectural variants tested. Each variant differs from the standard MHCflurry 0.9.1 predictor as indicated.

| MHCflurry variant | Description |
| --- | --- |
| *MHCflurry 0.9.1 (standard predictor)* | Two locally connected layers and a regularized fully connected layer of size 32 trained on quantitative and qualitative affinity measurements in IEDB |
| *quantitative* | Trained only on quantitative IEDB measurements |
| *0local-noL1* | No locally connected layers, no regularization on fully connected layer |
| *noL1* | No regularization on fully connected layer |
| *0local* | No locally connected layers |
| *1local* | One locally connected layer |
| *dense16* | Fully connected layer size of 16 |
| *dense64* | Fully connected layer size of 64 |

The HPV benchmark dataset consists of 194 affinity measurements across seven alleles. Peptides derived from the E6 and E7 proteins of HPV16 were assayed using a cell-based competitive binding assay [22, 23] (Supplemental Methods). We assessed accuracy on this benchmark using three well-known metrics, area under the receiver operator characteristic curve (AUC), F1, and Kendall rank correlation coefficient (Kendall’s tau). The AUC score estimates the probability that a strong binding peptide (measured affinity 500 nM or less) will have a stronger predicted affinity than a weak- or non-binding peptide (measured affinity greater than 500 nM). The F1 score summarizes a predictor’s precision and recall at classifying peptides as having affinity less or greater than 500 nM. The Kendall tau score measures the correlation in rank when peptides are sorted by measured or predicted affinity; it uses no cutoff and assesses agreement across the affinity spectrum.

## Results

MHCflurry 0.9.1 exhibited a modest improvement in accuracy over NetMHC 4.0 and NetMHCpan 3.0 in the ABELIN mass spec benchmark, outperforming NetMHC on 14/14 alleles tested and NetMHCpan on 12/14 alleles (Figure 2a). On average across alleles, MHCflurry showed a 10.9% (range 3.3 – 25.3) and 3.3% (-0.6 – +10.0) higher PPV than NetMHC and NetMHCpan, respectively.

**Figure 2:**
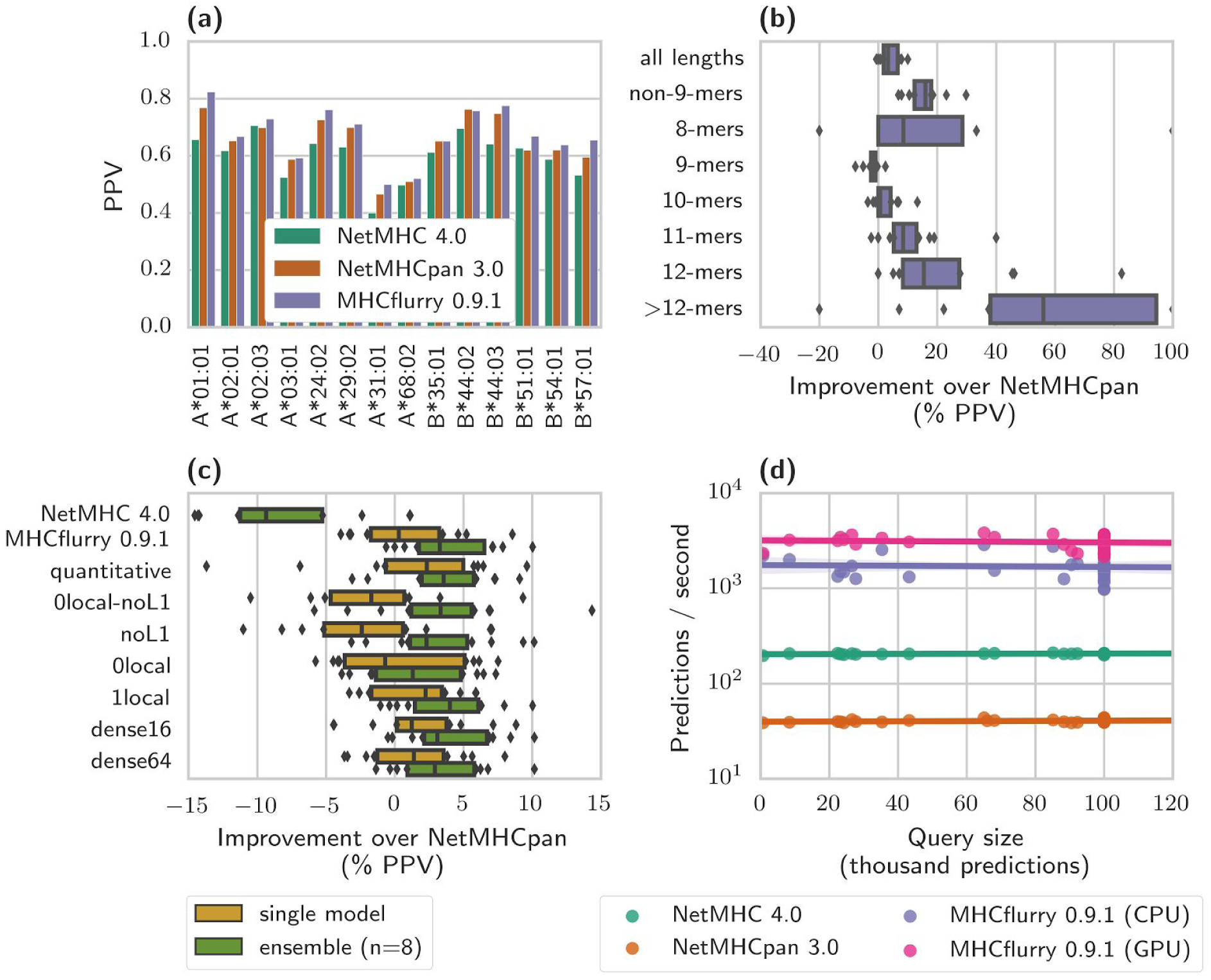
ABELIN mass-spec benchmark. **(a)** Positive predictive value (PPV) of NetMHC, NetMHCpan, and MHCflurry for each allele in the benchmark. **(b)** MHCflurry accuracy relative to NetMHCpan aggregated across alleles and split by peptide length. The median line is indicated, boxes show the quartiles, and points indicate individual alleles outside the interquartile region. The >12-mers category includes peptides of length 13, 14, and 15. **(c)** Relative accuracy of NetMHC, MHCflurry, and several variants of the MHCflurry architecture aggregated across alleles: *quantitative* is the standard architecture trained only on quantitative measurements in IEDB; *noL1* is an architecture with no L1 regularization on the fully connected layer; *0local* and *1local* indicate architectures with zero or one locally connected layers instead of two as in the standard MHCflurry architecture; *dense16* and *dense64* are architectures with a fully connected layer size of 16 or 64 instead of 32. Bars and points are as in (b). **(d)** Prediction speed. Indicated separately are timing measurements for MHCflurry when using only the CPU and when an NVIDIA Tesla K80 graphics processing unit (GPU) is available.

The accuracy advantage of MHCflurry over the NetMHC tools was due to better performance on non-9-mer peptides, which offset slightly lower accuracy on 9-mers (Figure 2b). On non-9-mers (i.e. peptides of lengths 8 and 10-15), MHCflurry outperformed both NetMHC and NetMHCpan on 14/14 alleles, with a median 38.4% (8.6 – 72.1) and 16.0% (range 6.9 – 29.9) improvement in PPV compared to NetMHC and NetMHCpan, respectively. On 9-mer peptides, MHCflurry underperformed NetMHCpan on 13/14 alleles, with a median difference in PPV of −1.7% (−7.7 – +2.4); relative to NetMHC the deficit was −1.0% (−3.0 – +5.0; underperforming on 10/14 alleles).

The architectural variants of MHCflurry tested showed overall similar performance, with all ensembles outperforming NetMHCpan in terms of median improvement in PPV across the 14 alleles in the ABELIN benchmark (Figure 2c). Similarly to the standard predictor, the architectures variants also showed a consistent advantage on non-9-mer peptides (Figure S1). The best performing architecture overall was the *1local* variant that incorporated a single locally connected layer instead of the two layers used in the standard MHCflurry 0.9.1 predictor. This variant obtained a median 4.0% (-1.0 – 10.0) higher PPV than NetMHCpan across alleles. Two other variants narrowly outperformed the standard predictor as well: the *quantitative* ensemble that was trained on only quantitative (not qualitative) IEDB data, and the *0local-noL1* predictor which used no locally connected layers and no regularization. While these three architectures outperformed the standard predictor in terms of median improvement in PPV, they also showed lower accuracy on their worst-performing alleles. The worst-performing allele relative to NetMHCpan for the standard MHCflurry 0.9.1 architecture was HLA-B*44:02, with a 0.6% lower PPV than NetMHCpan. For the *1local*, *0local-noL1*, and *quantitative* variants the worst performing alleles scored 1.0% (HLA-B*35:01), 5.8% (HLA-A*02:01), and 3.1% (HLA-B*35:01) lower PPV than NetMHCpan, respectively. This suggests that, while there is likely room for improvement, the MHCflurry 0.9.1 architecture and training strategy is a reasonable compromise between median and minimum performance across alleles.

Ensembles showed a consistent improvement in accuracy over single models, although even a single MHCflurry 0.9.1 model was on par with the NetMHCpan ensemble (median improvement=0.3%, range=-3.9 – +8.6). Ensembles also appeared to smooth out some of the difference between the architecture variants. For example, the *noL1* ensemble performed respectably (median=2.3%, range=-3.1 – 10.1) but a single *noL1* model performed much worse than the other models (median=-2.4%, range=-11.0 – +7.1), consistent with the idea that both regularization and ensembles can mitigate overfitting.

On the HPV benchmark, MHCflurry narrowly outperformed the other predictors in terms of AUC (MHCflurry=0.74, NetMHC=0.66, NetMHCpan=0.72) and F1 (MHCflurry=0.21, NetMHC=0.21, NetMHCpan=0.15). The NetMHCpan predictors outperformed MHCflurry in Kendall rank correlation (MHCflurry=0.15, NetMHC=0.17, NetMHCpan=0.19; Figure 3).

**Figure 3:**
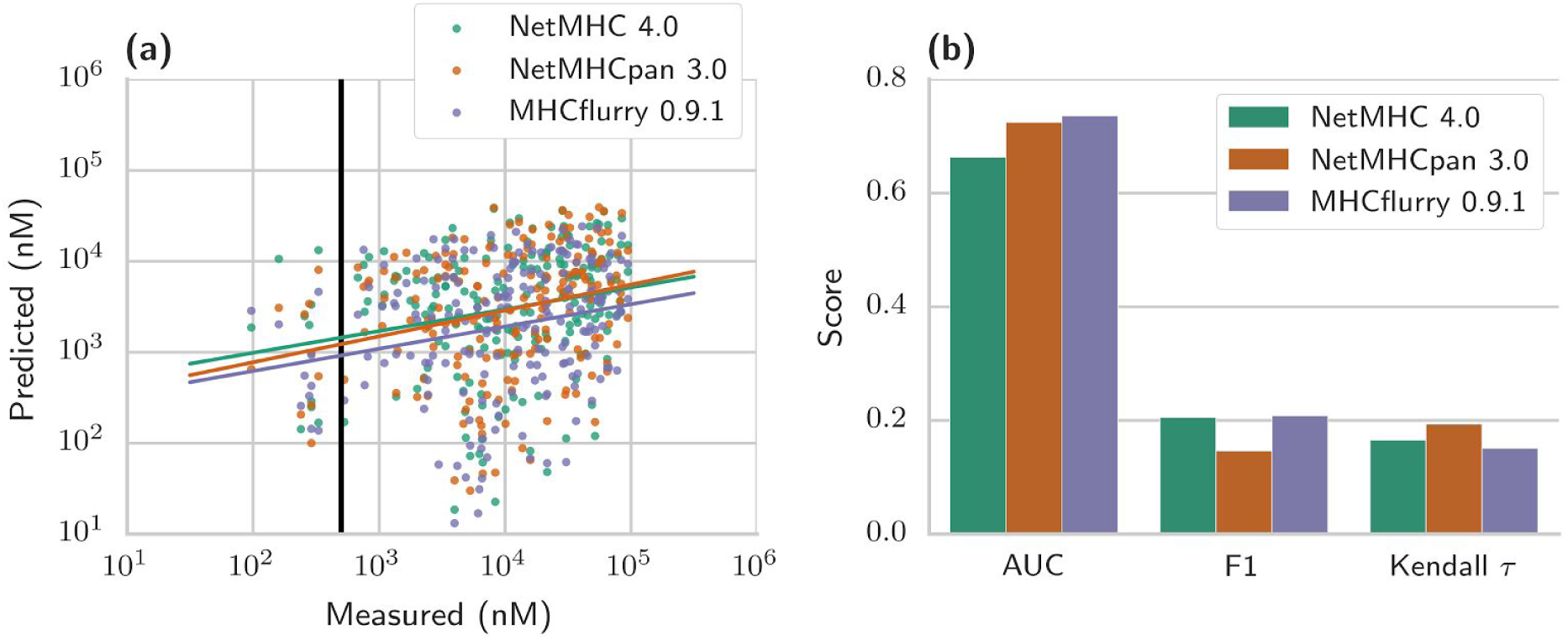
HPV affinity measurement benchmark. **(a)** Predictions for the three predictors on the HPV affinity dataset. The 500 nM threshold used in calculation of area under the receiver operating characteristic curve (AUC) and F1 scores is indicated. **(b)** AUC, F1, and Kendall rank correlation coefficient on the HPV dataset.

MHCflurry was substantially faster than the other predictors (Figure 2d). Using only the CPU, MHCflurry was on average 8.6X faster than NetMHC and 44X faster than NetMHCpan. Use of a GPU improved MHCflurry performance by about 75%, or 15X and 77X faster than NetMHC and NetMHCpan, respectively.

## Discussion

Here, we introduce a package of single-allele Class I MHC affinity predictors that achieves performance competitive with the well-known NetMHC and NetMHCpan tools. Our predictor, MHCflurry, supports several improvements over the neural network architecture and training approach used by NetMHC, including explicit support for variable length peptides, locally connected hidden layers, a regularized dense hidden layer, and incorporation of qualitative training data.

On the ABELIN mass-spec benchmark, MHCflurry outperformed the NetMHC tools overall and in particular on non-9mer peptides, suggesting that the peptide representation introduced here is a useful approach for training feedforward neural networks on variable-length MHC I peptide ligands. The non-9-mer accuracy improvement is interesting given that the IEDB training dataset is highly biased toward 9-mer peptides and contains very few peptides of length greater than 11 (1.4% of total). The case of 12-mer peptides is representative. In the ABELIN benchmark, the allele with the most 12-mer peptides is A*68:02, with 163 mass-spec identified 12-mers out of 1,986 total detected peptides. On this allele, the PPV scores were 0.20, 0.11, and 0.13 for MHCflurry, NetMHC, and NetMHCpan, respectively, suggesting that, while no predictor performs well in this context, MHCflurry learned something the other predictors did not. In the IEDB training data, there were only nine 12-mer peptides with affinity measurements for this allele, and all except one (TLVGLAIGLVLL with 542 nM affinity) had non-binder affinities.

This suggests that the MHCflurry models generalized to 12-mer prediction for this allele by learning from peptides of other lengths.

In contrast, MHCflurry narrowly underperformed the NetMHC tools on predictions for 9-mers, with a median 1.7% lower PPV than NetMHCpan across alleles on 9-mers. Addressing this deficit is important future work. As one possible explanation, we note that, unlike the NetMHC predictors, the MHCflurry training strategy performs no model selection; models for all alleles use an identical architecture. Model selection may be required for the last several percent in accuracy on 9-mers, a class of peptides for which the NetMHC tools are expected to be extremely well tuned.

MHCflurry has other important limitations. As a single allele predictor, MHCflurry cannot be expected to perform well on alleles with little training data in IEDB. NetMHCpan remains the best tool for such alleles. Furthermore, as only limited validation has been performed on MHCflurry models at this point, for clinical and other sensitive applications we recommend comparing MHCflurry predictions to existing predictors. In an effort to ease such comparisons, our group has developed the mhctools package (https://github.com/hammerlab/mhctools, manuscript in preparation), which provides a standard interface to running MHCflurry as well as popular binding predictors from other groups, including NetMHC and NetMHCpan.

MHCflurry’s prediction speed and convenient implementation may make it especially attractive for high-throughput epitope discovery efforts, such as neoantigen identification from high throughput sequencing of tumors. MHCflurry running on a CPU achieves nearly an order of magnitude speed improvement over NetMHC; this figure is greater still with respect to NetMHCpan or when using a GPU. Additionally, MHCflurry is readily integrated into scientific workflows as it may be installed using standard Python package infrastructure and includes both a command-line and a Python library API.

## Conclusion

MHCflurry is an open source Python package implementing MHC I affinity prediction for 101 alleles. On two benchmarks, it achieved accuracy competitive with the closed-source NetMHC and NetMHCpan tools and ran significantly faster. In contrast to these tools, MHCflurry supports training predictors on new datasets, is installed using standard package management infrastructure, provides a Python library API in addition to a command-line interface, includes automated unit tests and adheres to other software development best practices, and is distributed under a license that enables all users, including commercial entities, to use and improve the software free of charge. Epitope discovery efforts may consider integrating MHCflurry into their pipelines.

## Declarations

### Ethics approval and consent to participate

Not applicable.

### Consent for publication

Not applicable.

### Availability of data and material

The MHCflurry source code and training workflow, data, and models are available at https://github.com/hammerlab/mhcflurry. The TRAIN and ABELIN datasets (including predictions from all tools) may be downloaded at the following addresses:

TRAIN: http://github.com/hammerlab/mhcflurry/releases/download/0.9.1/data_curated.tar.bz2

ABELIN: http://github.com/hammerlab/mhcflurry/releases/download/0.9.1/abelin_peptides.all_predictions.csv.bz2

The HPV dataset includes unpublished affinity measurements (Hoppe, Bonsack et al., manuscript in preparation) and are available upon request from A. Riemer to not-for-profit enterprises for research purposes only.

### Authors' contributions

T.O. and A. Rubinsteyn developed MHCflurry. T.O. benchmarked the software and wrote the manuscript. M.B. and A. Riemer performed the HPV peptide binding experiments and advised on benchmarking approaches. J.H. supervised the project. All authors critically reviewed the manuscript.

## Acknowledgements

We thank Mike Rooney and Jenn Abelin for helpful discussions on binding predictor evaluation.

## Supplemental methods

### Competitive binding assay (HPV benchmark)

The binding affinity of HPV16 E6 and E7 derived peptides to selected HLA class I molecules was tested in competition-based binding assays as described in [22, 23]. Briefly, test peptides in 1:2 serial dilutions (final concentrations from 100 – 0.78 μM) compete with 150mM fluorescein-labeled reference peptide with a known high affinity for binding to the HLA class I molecule of interest on B-LCL cells, which were previously stripped from natural bound peptides and β2-microglobulin using ice cold citric acid buffer. After stripping, the cells were washed with culture medium and dissolved in culture medium containing 2μg/mL β2-microglobulin (MP Biomedicals) to reconstitute the HLA class I complex. B-LCL cells were diluted to 6x10^4^ cells/100μl per test peptide concentration and pipetted to a well-plate containing the mixes of test and reference peptide. After 24h incubation at 4°C the cells were washed, fixed in 1% PFA and suspended in 0.5% BSA in 1x PBS. The mean fluorescence intensity *Fmix* at each test peptide concentration was measured by flow cytometry (Accuri C6, BD Biosciences). The binding of each test peptide was calculated as the percent inhibition of reference peptide binding relative to the minimal response (without reference; *Fmin*) and the maximal response (reference only; *Fmax*) as:

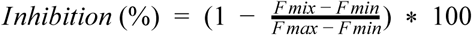

The binding affinity of the test peptide was determined by non-linear regression analysis as the concentration that inhibits 50% binding of the fluorescein-labeled reference peptide (IC50).

Peptides with an IC50 above 100μM were defined as non-binders. For confirmation and statistical significance the assay was performed at least three times for binders and twice for non-binders.

### Supplemental tables and figures

**Table S1:** Training and validation dataset sizes by MHC allele.

|  | **TRAIN affinity measurements**<br>**(% quantitative)** | **HPV affinity**<br>**measurements** | **ABELIN mass**<br>**spec ligands** |
| --- | --- | --- | --- |
| **HLA-A\*01:01** | 6,206 (67%) | 8 | 1,244 |
| **HLA-A\*02:01** | 26,254 (64%) | 34 | 2,574 |
| **HLA-A\*02:03** | 8,505 (87%) | 0 | 1,830 |
| **HLA-A\*03:01** | 9,634 (70%) | 10 | 1,662 |
| **HLA-A\*11:01** | 8,598 (70%) | 26 | 0 |
| **HLA-A\*24:02** | 5,851 (50%) | 46 | 2,206 |
| **HLA-A\*29:02** | 2,736 (87%) | 0 | 868 |
| **HLA-A\*31:01** | 6,278 (86%) | 0 | 1,229 |
| **HLA-A\*68:02** | 7,744 (93%) | 0 | 1,986 |
| **HLA-B\*07:02** | 6,935 (60%) | 20 | 0 |
| **HLA-B\*15:01** | 6,536 (63%) | 50 | 0 |
| **HLA-B\*35:01** | 4,266 (47%) | 0 | 894 |
| **HLA-B\*44:02** | 2,230 (88%) | 0 | 1,073 |
| **HLA-B\*44:03** | 1,491 (82%) | 0 | 912 |
| **HLA-B\*51:01** | 2,950 (86%) | 0 | 1,415 |
| **HLA-B\*54:01** | 1,234 (78%) | 0 | 1,212 |
| **HLA-B\*57:01** | 2,912 (90%) | 0 | 1,256 |
| **84 other alleles** | 125,237 (78%) | 0 | 0 |
| **All 101 alleles** | 235,597 (75%) | 194 | 20,361 |

**Figure S1:**
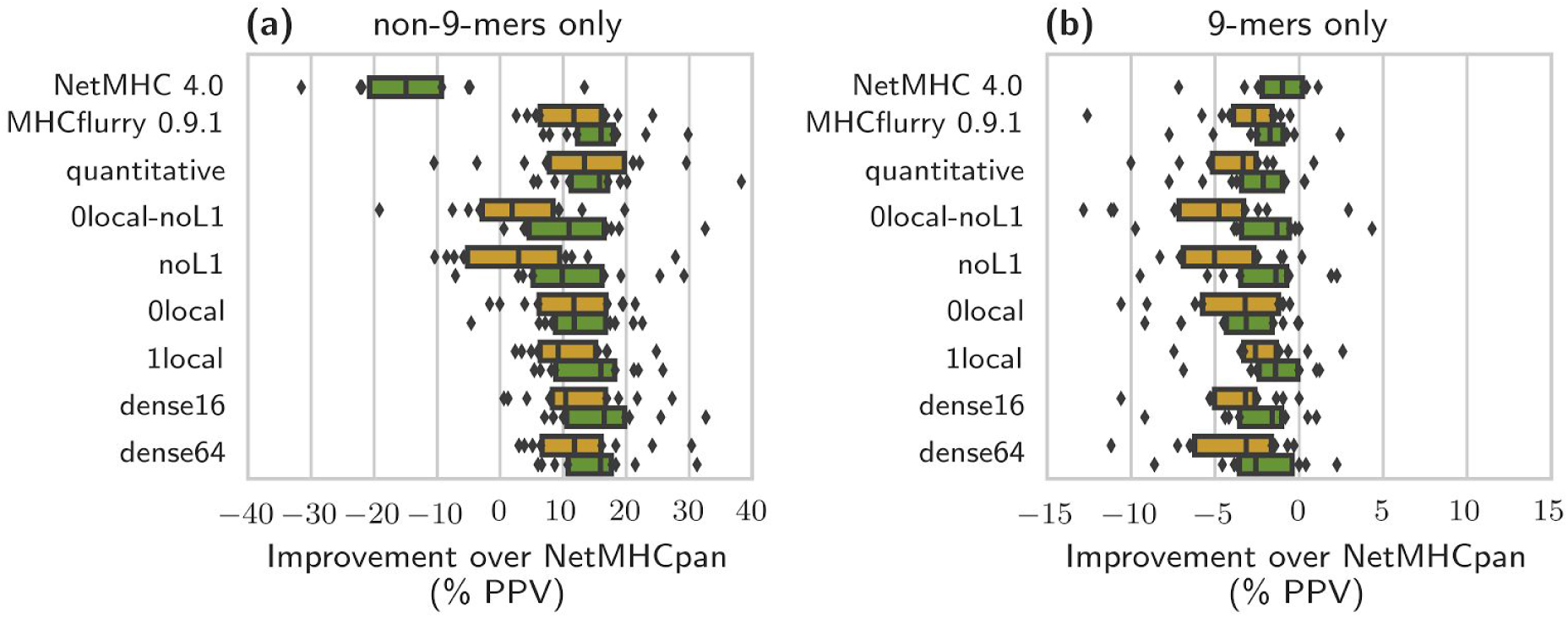
ABELIN mass-spec benchmark for NetMHC and several variants of the MHCflurry architecture for non-9-mer peptides (a) and 9-mer peptides (b). The bars and points are as in main text Figure 2(c).

